# Direction-selective retinal ganglion cells encode motion direction uniformly, despite having discretely distributed cardinal preferences

**DOI:** 10.1101/2025.07.23.666360

**Authors:** Carlo Paris, Félix Hubert, Felix Franke, Olivier Marre, Matthew Chalk, Ulisse Ferrari

## Abstract

On-Off direction-selective retinal ganglion cells (DS RGCs) exhibit broad tuning curves, responding robustly to motion aligned with or near their preferred direction. These cells comprise four major subtypes, each tuned to motion along one of the four cardinal axes: nasal, superior, temporal, or inferior. However, natural stimuli can move in any direction, and it remains unclear whether intermediate directions are encoded less effectively, or whether this cardinal organization nevertheless supports uniform direction encoding. Here, we combined previous electrophysiological recordings with an information-theoretic measure, the Stimulus Specific Information, to estimate the directional sensitivity of small populations of recorded neurons. This analysis revealed that DS RGC populations are uniformly sensitive across all directions of motion. We then asked whether the observed homogeneous sensitivity was a consequence of DS RGCs maximizing the average stimulus information. Simulations with artificially modified tuning curve widths revealed that DS cells prioritize avoiding pronounced drops in sensitivity over maximizing the average transmitted Information. Maximizing this minimal sensitivity may therefore be a principle to understand the organization of sensory systems.

## I. INTRODUCTION

Direction selectivity is one of the fundamental tasks of visual perception, evidenced by the presence of direction-selective (DS) cells in the visual system of a wide variety of species ranging from zebrafish larvae [1] to macaques [2]. DS cells respond robustly to motion aligned with or near their preferred direction, are among the most extensively studied types of cells, and have been observed in cat, ferret and macaque cortex [3–5], as well as salamander, mouse, rabbit and, recently, macaque, retina [6–8], to name a few. A striking difference between retinal and cortical DS cells is that, whereas the cortical cells exhibit a continuous spectrum of all preferred directions, DS retinal ganglion cells (RGCs) exhibit preferred directions that are restricted to a limited set of discrete axes. Given this difference, it remains unclear how functional organization of preferred directions impacts coding.

On-Off DS RGC respond strongly to both the onset and offset of a contrast change in their receptive fields if an object is moving along their preferred direction of motion [8, 9]. Furthermore, rabbit On-Off DS RGCs feature 4 preferred directions, roughly corresponding to cartesian axes: nasal, ventral, temporal and dorsal [10]. Intermediate directions of motion cause intermediate spiking with a slow monotonic decrease as the direction of motion moves away from the preferred one, resulting, in the rabbit retina, in broad tuning curves [13, 14]. Previous studies found a correspondence between these directions and the directions of apparent motion across the retina produced by contractions of the 4 rectus muscles, and suggested that these cells might control optokinetic nystagmus [10–12]. However, real-world visual stimuli can move in any direction, and from a coding perspective, it remains unclear whether motion along intermediate directions is poorly encoded by retinal DS cells.

Noting the small number of roughly equally spaced preferred directions, we inquire what the sensitivity to different stimuli looks like for a direction-selective system characterized by only four preferred directions. For example, do the large firing rates associated to stimuli aligned with the preferred directions lead to better encoding of those movements? In that case the system’s sensitivity would suffer a decrease for directions between the 4 preferred ones. Or, conversely, are intermediate directions better encoded than the preferred ones due to multiple On-Off DS RGCs of different types firing at intermediate capacity?

In order to estimate the system’s sensitivity to any direction of motion, we use information theory to link stimulus and neural response. Information theory [15] has a long history of application to neural systems and many studies successfully used measures such as the Mutual Information to estimate the encoding capacity of sensory neurons [16–18] (for review see [19]). However, Mutual Information is an average quantity and does not reflect sensitivity to individual directions of motion. To address this issue, here we use the Stimulus Specific Information (SSI) [20, 21], a decomposition of the Mutual Information which quantifies the contribution of individual stimuli.

A key result of this paper is that the tuning curve widths of On-Off DS RGCs maximize the minimum sensitivity, rather than maximizing the average sensitivity (i.e. the Mutual Information). To demonstrate this result, we applied an information-theoretic analysis to previous retinal recordings in rabbit [22, 23]. We begin by introducing and characterizing the dataset, in which, for quadruplets of DS RGCs - beach comprising one cell per type - we observe that sensitivity is approximately uniform across motion directions. Then, by artificially varying the widths of the tuning curves, we find that DS RGCs prioritize maximizing the minimum sensitivity across all possible directions of motion. In other words, the widths and preferred directions of the population are arranged such that the system avoids having any motion directions that are encoded significantly worse than others, thereby preventing the emergence of a kind of “blind spots”. Finally, we show that the cells converge on parameters that yield a “good enough” coding performance, rather than an optimal one, which may be physiologically too costly to achieve.

## II. RESULTS

### A. On-Off Direction Selective Retinal Ganglion Cells

We aim to quantify the sensitivity to motion in all possible directions of On-Off DS RGCs. We use previously recorded data collected from ex vivo New Zealand White rabbit retinas [22, 23]. Spiking activity was recorded from n=82 On-Off DS RGC, responding to bars moving in 36 equidistant directions, across 7 experiments (Methods). The cells show wide, bell-shaped tuning curves and can be typed according to their preferred directions, which are distributed around the 4 cardinal axes of the body - nasal, superior, temporal, and inferior (Figs. 1A,B).

**FIG. 1.**
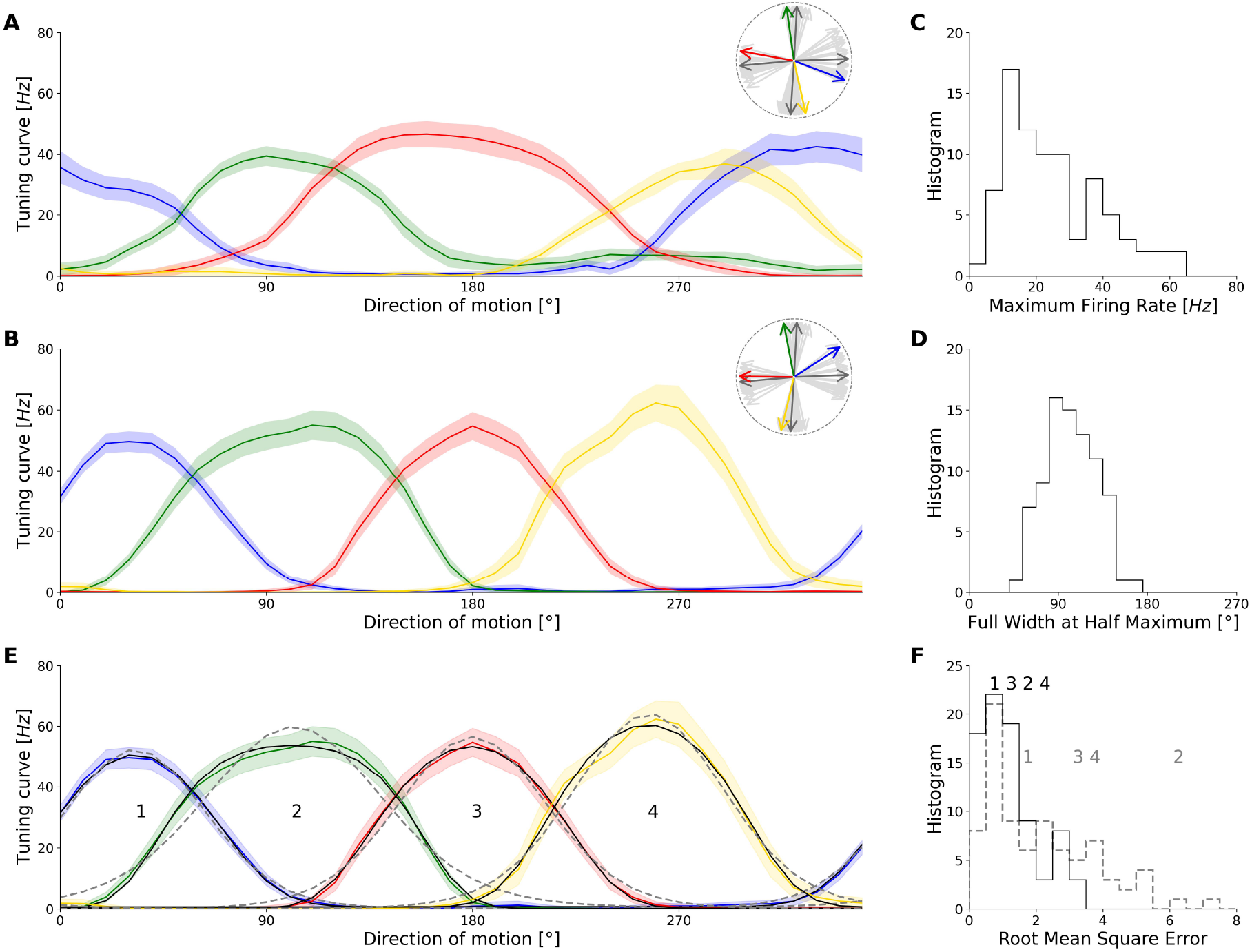
**A:** Example quadruplet of On-Off Direction Selective Retinal Ganglion Cells’ tuning curves. Each cell belongs to one of four types identified through their preferred directions (nasal - blue; superior - green; temporal - red; inferior - yellow). Solid lines and colored strips show the firing rates’ average and standard deviation, respectively, over 100 trials for each direction of motion. Inset: distribution of all the cell’s preferred directions (light gray) and the average preferred direction for each type (dark gray) compared to the preferred directions belonging to the cells in plot (coloured). **B:** Same as A, but for another quadruplet. **C:** Distribution of maximum firing rates over N=82 cells. *mean*±*std* = 40±22 **D:** Distribution of tuning curves’ widths, calculated as Full Widths at Half Maximum. *mean*±*std* = 103°±26°. **E:** Comparison between regular Von Mises (solid black line, rVM) and Flat-Topped Von Mises (dashed black line, FTVM). The commonly used rVM fit fails to reproduce the tuning curves’ curvatures near the maximum and the baseline firing rates. On the other hand, the FTVM manages to do so remarkably well, best exemplified by tuning curve 2. **F:** Comparison of Root Mean Square Error distributions between data and fitted tuning curves for the rVM (cyan) and FTVM (black) fitting functions. The numbers 1-4 reference the tuning curves from panel E, and show the RMSE reduction in switching from a rVM to a FTVM fit.

Cells exhibit a wide distribution of maximal firing rates, ranging from 10*Hz* to 80*Hz*, with most cells peaking at 20*Hz* (40±22*Hz*, Fig.1C). Notwithstanding the diversity of maximal firing rates, the cells’s tuning curves, when normalized, resemble one another, as evidenced by the fact that most of the cells have full widths at half maxima (FWHM) close to 100° (103°± 26°, Fig.1D). Additionally, the tuning curves overlap noticeably for orthogonal preferred directions (Figs.1A,B and D), and because of this we expect the sensitivity of a population of DS RGCs to behave very differently from the sensitivity associated with individual cells.

### B. The sensitivity of small populations of On-Off DS RGCs is homogenous across directions

Real-world stimuli move in all directions, yet DS RGCs exhibit preferred directions which align along only four axes. This raises the question of whether populations of these cells maintain uniform sensitivity across all directions of motion, or whether the encoding of certain directions is significantly compromised due to this specific distribution of preferred directions. To test this, we consider combinations of 4 cells - one cell for each type - as units of computation. This approach is motivated by the fact that On-Off DS RGCs tile the retina such that receptive fields of cells with the same preferred direction exhibit minimal overlap, whereas receptive fields of different types overlap randomly [24]. Consequently, at any given point on the retina, a small stimulus is processed by only four cells, one from each type.

We form quadruplets by exhaustively combining tuning curves of different types within individual experiments. For each quadruplet we calculate its stimulus sensitivity using the *Stimulus specific information* (Fig. 2A,B, Methods and [20, 21]. We choose this quantity because it measures the reduction of stimulus uncertainty (entropy) after observing the cells’ response and for this data it aligns with decoding performance better than other quantities (Suppl. Sect. A). Additionally, its average over the stimulus ensemble yields the Mutual Information between stimulus and response [20], and this allows us to to compare individual stimulus sensitivites to overall performance.

**FIG. 2.**
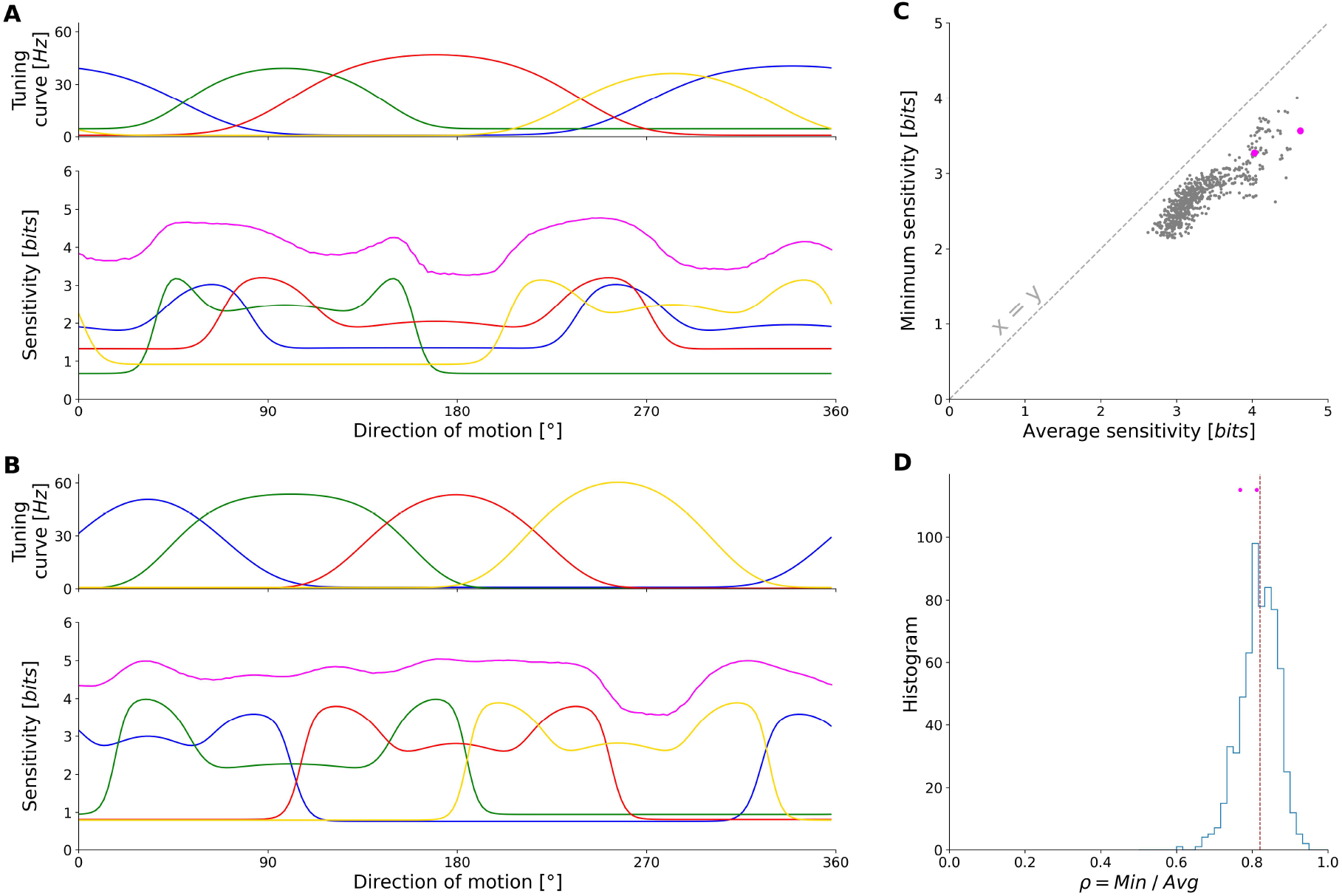
**A:** Upper - fits for tuning curves from Fig. 1A; lower - associated individual sensitivity (non-magenta colors) and population sensitivity (magenta). **B:** Analogous to A, but for the quadruplet in Fig. 1B. **C:** Average sensitivity versus Minimum sensitivity for each randomly formed quadruplet (N=648). Magenta points indicate the sensitivities of quadruplets from A andB. Equality line identifies cases where the sensitivity is constant, with *Min*(*SSI*) = *Avg*(*SSI*). **D:** Distribution of quadruplets’ minimum sensitivities relative to the respective averages (dotted line shows the median). 439 out of 648 quadruplets (67.7%) show a minimum sensitivity greater than 80% of the average.

As previously observed, the stimulus sensitivity of individual cells is large around their preferred directions (Fig. 2A,B), with typical peaks corresponding to the tuning curves’ flanks [21]. On the other hand, quadruplet sensitivity expresses the stimulus information encoded by a group of four cells, and in the examples presented it appears fairly homogeneous (Fig. 2A,B magenta lines). Upon obtaining the sensitivities for all quadruplets, we notice that the minimum of the sensitivity often results close to the average value (Fig. 2C) - the mutual information. This indicates that the sensitivities of the quadruplets oscillate conservatively around the average. More than half of the quadruplets show a minimum sensitivity value within 20% of the average (Fig. 2D). In other words, the difference between any given quadruplet’s Mutual Information and minimum sensitivity lies between 0.5 and 1 bit of information.

A flat sensitivity is achieved through a balance between maximal spiking of a cell for stimuli moving near its preferred direction, on the one hand, and multiple cells spiking at intermediate levels when the stimulus lies between preferred directions, on the other hand. Populations of four DS RGCs thus exhibit uniform sensitivity to all possible directions of motion. This property should result from the specific shape and width of the tuning curves. Therefore, in the next section we address how overall stimulus sensitivity - and its directional uniformity - depends on the shape of individual tuning curves.

### C. Widths of On-Off DS RGC tuning curves are tuned to maximize sensitivity minima

The minimum of the sensitivity is associated with the least informative, or most poorly encoded, stimulus. Increasing this value would be advantageous for the system as it would simultaneously eliminate a potentially fatal “blind spot” and heighten the overall average. On the other hand, the average sensitivity corresponds to the Mutual Information *MI* [20], a measure of the average amount of information the system can transmit. Previous works argued that sensory systems evolved to maximize this quantity under energetic constraints, a hypothesis known as the Efficient coding principle [25–27]. However, as maximizing the average or overall sensitivity might leave the aforementioned “blind spots”. Given the small differences between quadruplets’ Mutual Information and minimum sensitivities (Fig.2D) we hypothesized that the cells have tuning curve widths which yield sensitivities as constant as possible along the stimulus ensemble. To test this hypothesis, we simulate how each quadruplet’s sensitivity changes upon modifying the constituent tuning curve widths (Methods) and focus on minimum sensitivity and Mutual Information.

For each modified width of a given quadruplet, we calculate the quadruplet’s minimum sensitivity and the corresponding Mutual Information (average sensitivity) (Fig.3A). We can plot the last 2 quantities against the associated average tuning curve widths whereupon we obtain curves which, as expected, drop towards 0 at the extremes. Indeed, as the tuning curves become wider and wider, the cells’ activities become increasingly uniform across directions of motion, ultimately losing direction selectivity altogether (Fig.3A right). On the other hand, as the tuning curves become narrower they tend towards peaked functions centered in the system’s 4 preferred directions of motion. In those directions, the encoding is perfect (so the sensitivity equals the stimulus entropy), but other directions are completely overlooked (leading to null sensitivity). In that case the minimum of the sensitivity goes to zero (Fig.3A left).

By performing simulations of systematic compression and broadening of the tuning curve widths, we observe two different behaviors for minimum sensitivity and Mutual Information. The minimum sensitivity can present either 1 or 2 clearly visible local maxima (exemplified in Fig.3B, blue lines), located close to either the empirical or relatively much narrower widths. Consistently, when studying all the *N* = 648 possible quadruplets, the histogram of the widths corresponding to the optimal minimum sensitivity is bimodal, with two peaks centered at values similar to or smaller than the empirical widths (blue histogram in Fig.3C). Although the globally optimal average tuning curve width for a given quadruplet may be much narrower than the empirical one, the resulting difference in minimum sensitivity is typically small. In fact, more than half of the empirical quadruplets exhibit a loss of less than 0.2 *bits* relative to the globally optimal (i.e., highest) minimum sensitivity (blue histogram in Fig.3D; median = 0.184 *bits*). Overall, these results indicate that the widths of the empirical tuning curves are tuned to values that yield near-optimal minimum sensitivity for the quadruplets of cells.

In contrast, we observe a different outcome for the Mutual Information (average sensitivity). For most quadruplets, we observe a single maximum (Fig.3B, red lines), typically occurring at tuning curve widths much narrower than the empirical ones (Fig.3C, red histogram). Accordingly, the difference between empirical and optimal average sensitivity is often substantial, with only about 10%of quadruplets showing a loss smaller than 0.2 bits (Fig.3D, red histogram, median = 0.433 *bits*).

These results suggest that empirical quadruplets are tuned to maximize minimum sensitivity rather than Mutual Information. Nevertheless, the quadruplets seem to reach a near-optimal minimum sensitivity, rather than achieving the globally optimal configuration. Next, we wonder whether this solution represents a compromise by incurring a marginal loss in performance in order to avoid an optimization toward very narrow widths.

### D. Widths of On-Off DS RGC tuning curves are “good enough”

For about half of the quadruplets, the width that maximizes the minimum sensitivity is narrower than the empirical one (Fig.3C), and is comparable to the width that maximizes average sensitivity. Yet the empirical quadruplet exhibits much broader tuning curves while still incurring only minor losses of performance in terms of minimum sensitivity. Motivated by these findings, we test whether these broader, but “good enough”, widths could result from an optimization that starts from unspecific cells and gradually sharpens their tuning.

For each quadruplet, we start from an avatar system with very large widths, and progressively narrow them by crawling leftwards along the curves of either the minimum sensitivity or the Mutual Information (Fig.4A), stopping at the first local maximum of the curve (Methods). The example quadruplet in the top panel of Fig.3A illustrates a case where the “crawling” procedure converges to an optimal minimum sensitivity that, while globally sub-optimal, closely matches the empirical one (Fig.4A top). In contrast, for the quadruplet shown in the bottom panel of Fig.3A, the same procedure yields the globally optimal width, identical to the one found earlier (Fig.4A bottom). In both cases, optimization of the Mutual Information converges to a single, global maximum (Figs. 3A and 4A).

**FIG. 3.**
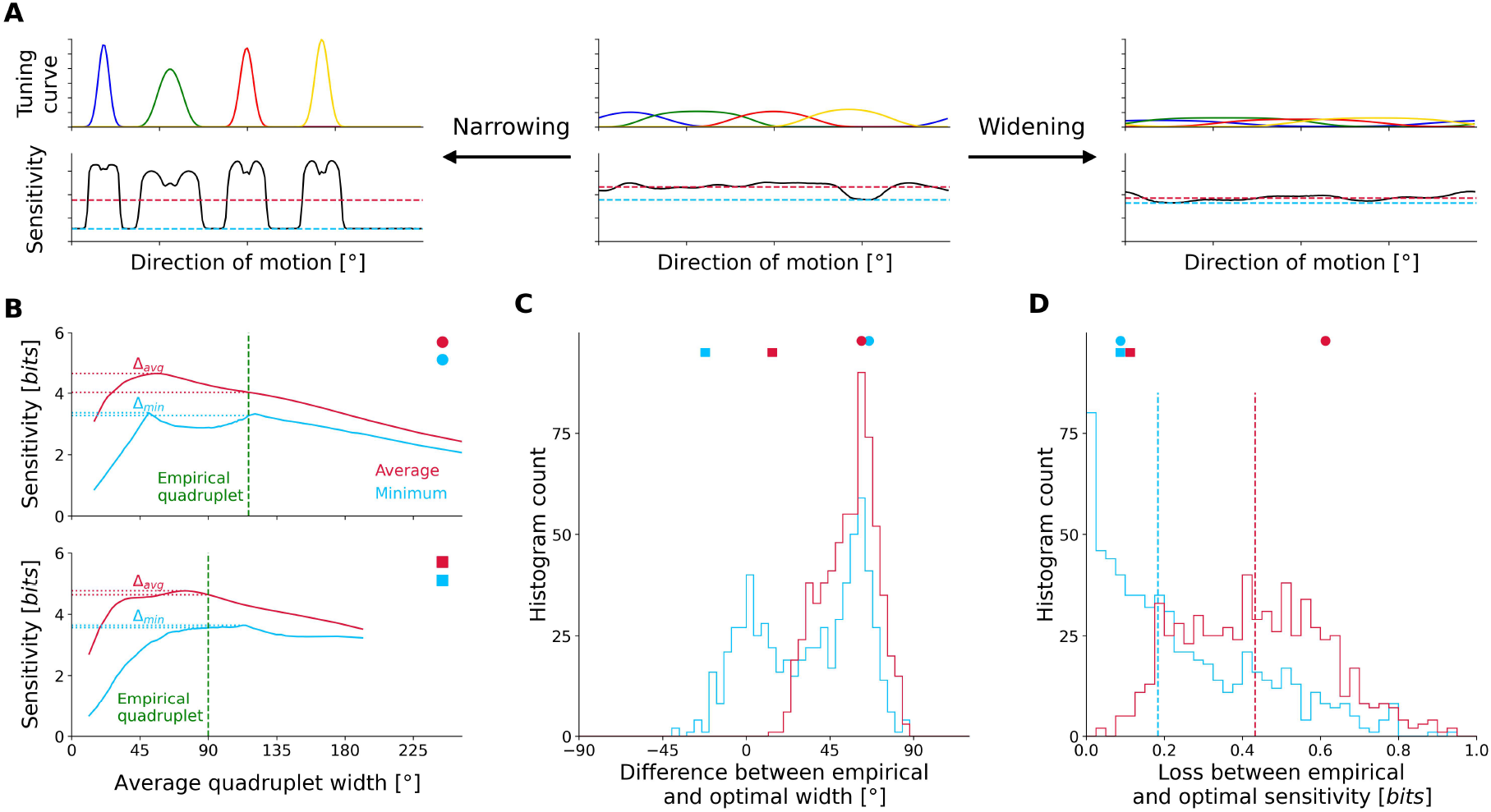
**A:** Simulation of various tuning curve width regimes. Starting from empirically obtained widths, quadruplets of “synthetic” avatars are generated by modifying the cells’ widths to smaller or larger values. **B:** Average (red) and minimum (blue) sensitivity as a function of the stretched average tuning curve width for the quadruplets from Fig.1A and B. The green line is the average Full Width at Half Maximum (FWHM) of the 4 empirical tuning curves forming this quadruplet. Δ*_avg_*is the difference between the maximum average sensitivity and the average sensitivity achieved by the empirical quadruplet (red). Δ*_min_*is analogous, but for the minimum sensitivity (blue). **C:** Histograms of differences between the average width of a given empirical quadruplet and the average width at which this quadruplet maximizes either the average (red) or the minimum sensitivity (blue). A positive difference means the empirical quadruplet is wider than its maximizing avatar. The circle and square symbols mark the bins where the quadruplets from panel B fall. **D:** Histograms of Δ*_avg_*(red) and Δ*_min_*(blue). These quantities express a loss with respect to the case in which the quadruplet maximizes either the average or the minimum. The dashed lines indicate the histograms’ median values. Note how the Δ*_min_*median is smaller than that of Δ*_avg_*, indicating that empirical quadruplets have an almost optimal minimum sensitivity.

**FIG. 4.**
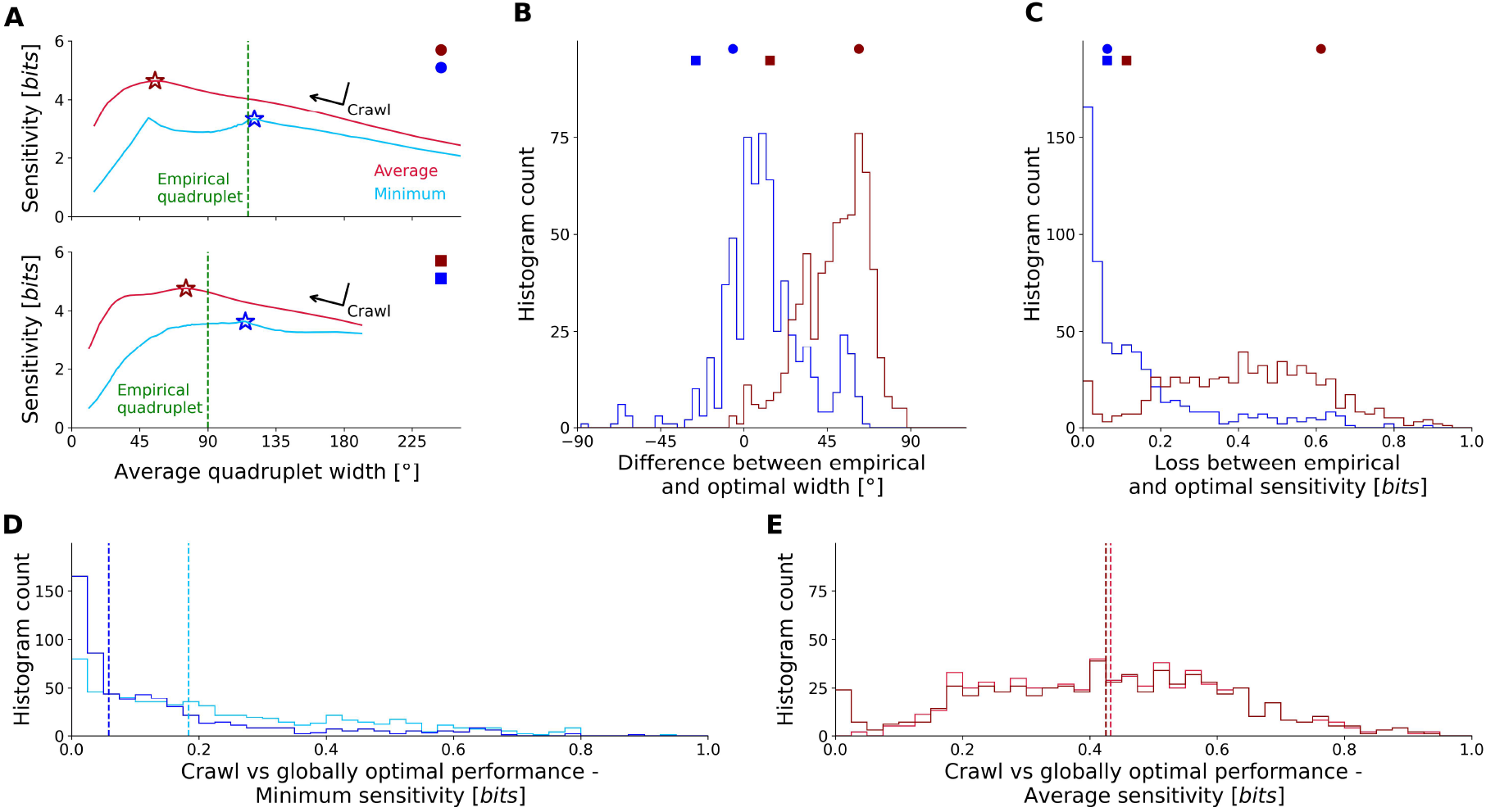
**A:** For each quadrulet, starting from large average tuning curve widths (far right) and crawling leftward, we find the first local maximum along either the minimum or average sensitivity curve. The darkred and darkblue stars mark the 2 resulting peaks. **B:** Similar to Fig.3C, histogram of the difference between their average width in the empirical regime and the quadruplet’s width yielded by the crawling procedure. The latter may or may not correspond to the globally optimal width. Circles and squares mark the quadruplets from panel A. **C:** Histogram of performance losses between the sensitivities in the empirical regime and the optimal sensitivities reached by the crawling procedure. **D:** Comparison between the performance losses estimated by finding the global maximum of the minimum sensitivity (Fig.3D, blue) versus the losses of minimum sensitivity estimated through the crawl procedure (panel D, darkblue). The dashed lines mark the distributions’ medians: dark blue 0.06*bits*, light blue 0.18*bits*. **E:** Analogous to panel D, but for the average sensitivity

An analysis including all possible quadruplet combinations confirms these results. Our crawling procedure for the minimum sensitivity produces widths that closely match the empirical ones (Fig.4B and C, blue histograms). Consistently, the performance losses with respect to the empirical width of the crawl optimum is smaller than those of the global one (Fig.4D). Conversely, applying the crawling procedure to the Mutual Information converges to the same optimal width identified previously (compare red histograms in Fig.3C and 4B), with correspondingly large performance losses. Regardless of possible underlying mechanisms or interpretations (see Discussion), our results indicate that empirical widths lie near a local optimum of minimum sensitivity - one that can be reached by progressively sharpening initially broad, directionally non-selective tuning curves.

## III. DISCUSSION

In summary, we investigated direction selective retinal ganglion cells (DS RGCs) from rabbit retinas to determine whether the discrete distribution of four equidistant preferred directions results in homogeneous stimulus sensitivity - and whether this organization may be justified by an optimization principle. To address this question, we quantified the sensitivity using a decomposition of the Mutual Information along the stimulus ensemble and found that randomly assorted quadruplets of On-Off DS RGCs exhibited uniform sensitivity across directions of motion. We then showed, through simulations across widths, that the empirical tuning curve widths place the cells in a regime that doesn’t optimize Mutual Information (average sensitivity), but instead yields minimum sensitivities close to the globally optimal ones. Despite this, the empirical widths were rarely globally optimal with respect to minimum sensitivity, suggesting that the cells operate in a “good enough” regime that results from an optimization that gradually sharpens the tuning curves (Fig.4).

We observed that empirical widths lie close to a local maximum of the minimum sensitivity, rather than the global maximum, which occurs at sharper tuning curves. From a developmental standpoint, although widths narrower than the empirical ones may yield higher minimum sensitivities (Fig.3C), achieving them may be unfavorable due to biological constraints. Biological widths are arrived at through a gradual development of asymmetric synaptic connections between Starburst amacrine cells and DS RGCs during the organism’s postnatal maturation [28, 29]. In rabbits and mice, the distribution of preferred directions and the tuning curve widths have been observed to develop after eye opening, with preferred directions aligning progressively with the canonical axes and tuning curves becoming narrower [30]. If synaptic asymmetry underlies direction selectivity, then an initially asymmetric connectivity would be associated with completely flat, directionally non-selective tuning curves. As development proceeds, the emergence of asymmetry would lead to reduced firing in the anti-preferred direction, accompanied by a gradual relative increase in response near the final preferred direction. Optimizing the tuning curve width beyond a certain level of performance likely incurs for the system some form of energy and wiring strength cost which may outweigh the benefits.

Although our study focuses on the role of tuning curve widths, other parameters may also contribute to achieving optimality. Coding performance depends on the spacing of preferred directions, with equal spacing enabling optimal information encoding, and deviations from this arrangement leading to reduced performance. Theunissen et al. [31] showed that sliding one tuning curve while keeping others fixed leads to a monotonic decrease in performance, reaching a minimum when the sliding tuning curve overlaps with one of its neighbors. Our results are compatible with these findings, as the vacancy created by the shifted tuning curve would significantly reduce sensitivity, thereby violating the Max-Min principle. Similarly, the placement and number of preferred directions also impacts coding, as shown in [22] using the same dataset as the present study. While perfectly spaced and more numerous preferred directions lead to lower decoding error, Fiscella et al. propose that the upgrade might not be conducive to a net improvement, considering the resource cost necessary to achieve it. Our results regarding uniform sensitivity for a system with cells selective to a discrete number of preferred stimuli align with Theunissen et al. [31], even though they worked on a completely different biological system (the cricket cercal system). Notably, both systems strive to maximize the minimum sensitivity, supporting this goal as a general encoding principle to understand the organization of sensory systems.

Technically, Theunissen et al. [31] measured sensitivity with a different decomposition of the Mutual Information. However, both their chosen measure and ours can be related to the stimulus dependent decoding error *σ_dec_*(*θ*) through the same functional form (see Appendix of [31] and our Suppl. A, and Protocol 2 of Butts and Goldman [21] for a more detailed description of how different information-based sensitivity measures compare to each other). We chose to work with a different measure - the Stimulus Specific Information [20, 21] - because it quantifies the residual uncertainty when decoding the stimulus from the neuronal responses (App. A), making it particularly well-suited for investigating how individual stimuli are represented. In our opinion, there is no single right decomposition of the Mutual Information, and the choice should depend either on interpretability or the specific system function one wishes to study, following criteria such as additivity [32], causality [20], coordinate invariance [33] and locality [34].

In the future it will be interesting to see how well the Max-Min sensitivity principle holds up in other systems. For instance, ON RGCs in the rabbit retina have been shown to be characterized by 3 roughly equidistant preferred directions [10, 12, 35]. We could wonder if also their tuning curve width is explainable through the same principle. Additionally, equivalent questions might be posed for cortical cells, such as DS cells in the primary visual cortex which share the same one-dimensional stimulus ensemble with their retinal counterparts, or even place cells whose stimulus ensemble is at least two-dimensional. Answers to these questions will pave the way to a more complete understanding of the role specific cell types play in perception, and the Max-Min principle can serve as a compass to guide that research.

## IV. METHODS

### Experimental data

The experimental protocols has been previously presented, see [22, 23] for more details. Briefly, the data was recorded from New Zealand albino rabbit retinas in the course of 7 experiments. Each retina was extracted and placed ganglion cell side down on a high-density multi-electrode array featuring 11011 electrodes [37]. The light stimulation was performed using a light bar moving in one of 36 possible equidistant directions, each repeated 100 times. Cell activity was monitored online in order to find electrodes recording On-Off Direction Selective cell activities. Spike sorting was done manually using the UltraMegaSort2000 [38]. The spike sorted cells were subsequently classified as oo-DSRGCs according to the Direction Selectivity Index (DSI) and the On-Off Index (OOI), respectively defined as:

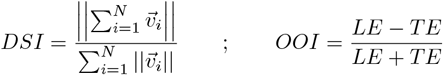

where 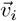 is the cell’s average firing rate in the *i_th_* direction, while *LE* and *TE* denote the responses in the cell’s preferred direction of motion to the leading and trailing edge when the bar is moving in the cell’s preferred direction of motion. Only those cells were kept which had *DSI* > 0.2 and *OOI* < 0.8. A total of *n* = 90 cells satisfied these requirements.

### Fitting the tuning curves

We model the tuning curves with a Flat-Topped Von Mises (FTVM) function, an extension of the regular Von Mises function (rVM) [23, 39, 40]]. Its functional form reads:

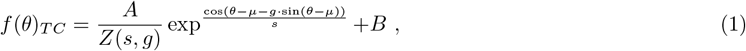

where *Z*(*s, g*) is a normalization constant.

The independent variables of the function are the directions of motion *θ*∈Θ. The remaining quantities are fitting parameters: µ is the cell’s preferred direction, *B* is the cell’s activity in the anti-preferred direction, *A* is the area above the baseline, *s* is the so-called concentration parameter which regulates the curve’s width (akin to the standard deviation in the 1-dimensional Gaussian case) and *g* is the flatness parameter. By setting *g* = 0, we can retrieve the regular Von Mises function. Reducing *g* results in a function which is more peaked than the *g* = 0 case, whereas increasing it flattens the top.

An additional selection of cells is done at this point by keeping only those cells for which the Root Mean Square Error of the FTVM fit was less than 5, and those whose FTVM fit featured a Full Width at Half Maximum less than 180°.

### Noise model

The conditional probability *p*(*r|θ*), and therefore the *I_SSI_* itself, depends on the choice of noise model. We assume a Poissonian noise model so that both the spiking average and variance for any stimulus *θ* are given by the corresponding value of the tuning curve *f_T C_*(*θ*), that is we assume the rate *λ* = *f_T C_*(*θ*). With these assumptions, the conditional probability reads

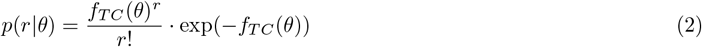

This is broadly supported by the data and at any rate seems to be an acceptable approximation (see Supplementary materials for further information).

Finally, unless stated otherwise, we assume conditional independence of the responses, meaning that for a quadruplet of cells responding to a stimulus *θ* ∈ Θ, the probability of responses (*r*_1_*, r*_2_*, r*_3_*, r*_4_) is equal to

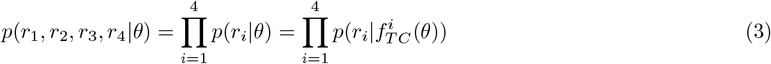

### A measure of sensitivity - Stimulus Specific Information

This paper studies how the width of On-Off Direction Selective Cell tuning curves impacts the sensitivity of this system to various directions of motion. We approach this problem from an information-theoretic perspective and use the Stimulus Specific Information (SSI) as a measure of sensitivity [20, 21], a decomposition of the Mutual Information over the stimulus ensemble:

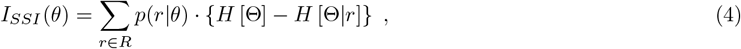

where *θ*∈ Θ is the light bar’s direction of motion, *R*∈ N_0_ is the cell response ensemble and *r* the spike count vector of the cells. *p*(*θ*) is the distribution of stimuli (here uniform), while *p*(*r θ*) is the probability that a given response *r* is elicited by a particular stimulus *θ* and *H*[·] are entropies. The term *H*[Θ]*− H*[Θ *r*] is the information gained about which stimulus was shown to the system upon learning that the system responded with *r*. The SSI is the average information gained by the system’s response, weighed by the conditional probability that a response was elicited by a particular stimulus. It bears an explicit dependence on the stimulus *θ* ∈ Θ and its average over the stimulus ensemble equals the Mutual Information between the two ensembles: *MI*[Θ; *R*]:

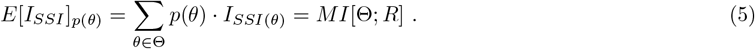

### Quadruplet formation

We consider as elementary computational units quadruplets of cells such that each cell belongs to one of the four subtypes according to their preferred direction of motion (superior, temporal, inferior, nasal). Such a choice follows from the fact that the receptive fields belonging to each of the 4 subtypes completely tile the retina independently from cells of the other 3 subtypes, with minimal overlap of same-subtype receptive fields [24, 41]. Consequently a small stimulus falling on a point of the entire visual field will most likely be directly processed by 4 On-Off direction selective retinal ganglion cells, namely 1 belonging to each of the 4 subtypes. Within every experiment, all possible combinations of 4 cells are taken while preserving the original sub-typing. Cells from different experiments are not mixed. The result is a total of *N* = 648 quadruplets.

### Tuning curve stretching simulation

Each cell’s FTVM fit has 5 parameters (Eq. 1), one of which is the concentration parameter *s* which controls the tuning curve’s width. A quadruplet of cells will correspondingly have 4 such concentration parameters and an average Full Width at Half Maximum.

In our simulation, in order to obtain a “new” quadruplet we multiply the 4 *s* parameters of the original empirical quadruplet by the same multiplier. All other (16) parameters are kept constant. Multiplying the original *s* parameters by a value lower or higher than 1 results in respectively narrower or wider tuning curves. Therefore by choosing an array of multipliers spanning values from 0.01 to 10 we are able to explore a wide variety of avatars of the original quadruplet, each differing from the others by their average Full Width at Half Maximum.

For each such quadruplet we calculate the associated Sensitivity as a function of the direction of motion, its minimum sensitivity min_*θ*∈Θ_*SSI*(*θ*) and average sensitivity *E*[*SSI*(*θ*)] (= *MI*[Θ; *R*]). Since the computational complexity of an exact calculation of the SSI increases explonentially with the number of cells, we implemented a Monte Carlo sampling method as described in [42]. Doing this for all multipliers (and thus all width regimes) yields the behavior of the minimum and average sensitivities as a function of the quadruplet’s average width.

## ACKNOWLEDGEMENTS

Felix Franke would like to thank the SNSF Eccellenza Grant (PCEFP3 187001), SNSF Spark Grant (CRSK-3 220987, F.F.) and SNSF Projects in Life Sciences Grant (310030 220209). Olivier Marre would like to thank the ERC Consolidator grant DEEPRETINA (101045253) and the Agence Nationale de la Recherche (ANR) for financial support (Chaire Industrielle MyopiaMaster ANR-22-CHIN-0006, ANR-20-CE37-0018-04-Shooting Star, ANR-22-CE37-0033 NUTRIACT, ANR-22-CE37-0016-01 PerBaCo and ANR-RetNet4EC. Matthew Chalk would like to thank the Agence Nationale de la Recherche (ANR) for financial support (ANR-RetNet4EC). Ulisse Ferrari acknowledges this work was done within the framework of the PostGenAI@Paris project with the reference ANR-23-IACL-0007, as well as having benefitted from financial support by the Agence Nationale de la Recherche (ANR) by grants NatNetNoise (ANR-21 CE37-0024), IHU FOReSIGHT (ANR-18-IAHU-01). Our lab is part of the DIM C-BRAINS, funded by the Conseil Régional d’Ile-de-France.

