## Supplementay S1 for "Direction-selective retinal ganglion cells encode motion direction uniformly, despite having discretely distributed cardinal preferences"

### SUPPLEMENTARY INFORMATION

#### A. The Stimulus Specific Information offers a good measure for the stimulus sensitivity

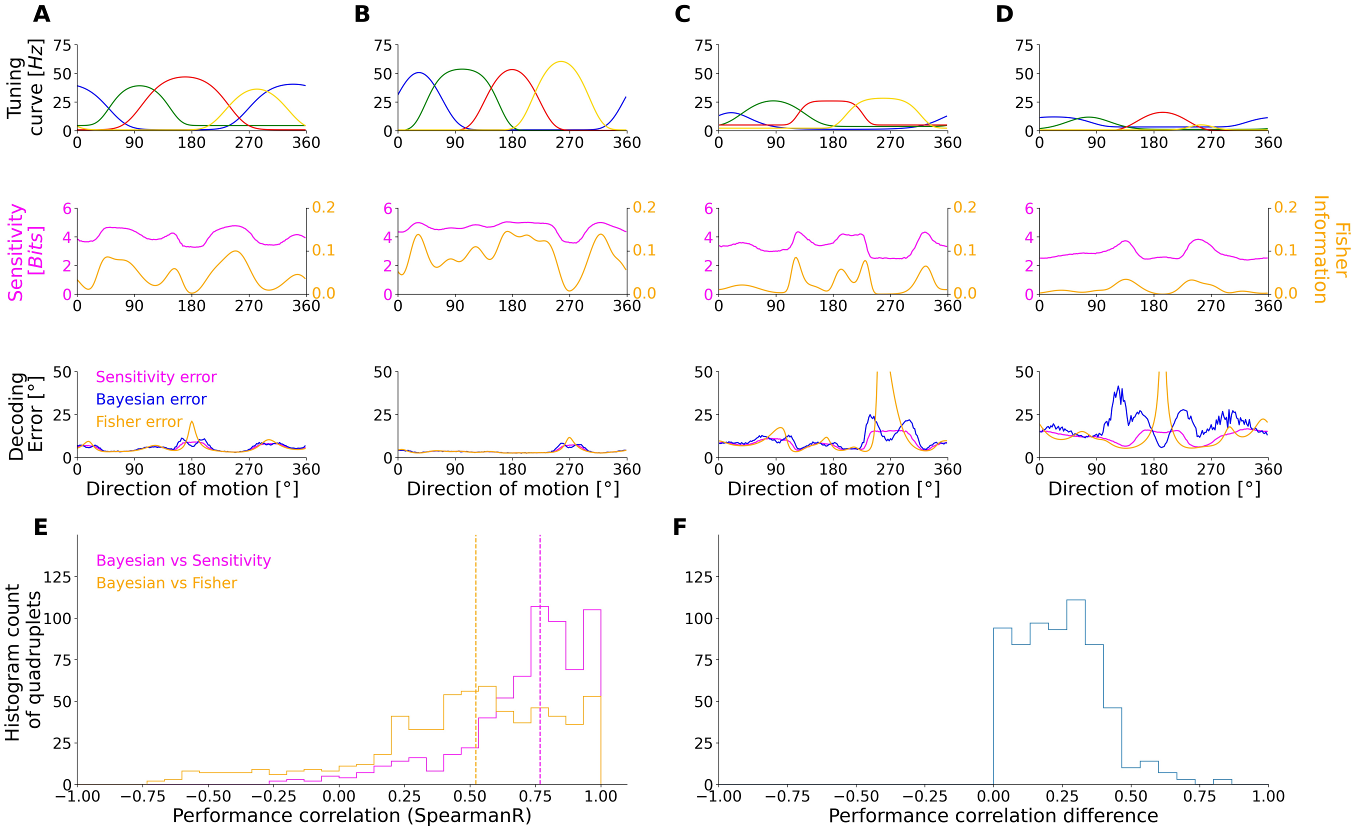


FIG. S1. **A:** upper - Example quadruplet of tuning curves from Fig.1A; middle - Population sensitivity of Fig.1A quadruplet in magenta and its Fisher information $FI(\theta)$ in orange; bottom - Decoding errors estimated from the sensitivity, Fisher Information or a Bayesian unbiased decoder. The sensitivity error is proportional to $e^{-SSI(\theta)}$ with the $SSI$ expressed in *nats*. The Fisher error is obtained from the Fisher information $FI(\theta)$ as $1/\sqrt{FI(\theta)}$, assuming an unbiased decoder with flat prior $p(\theta) = 1/|\Theta|$. Notice the Fisher information peaking around 180°, in correspondence to the red cell from the uppermost panel. This is due to the slopes of all 4 tuning curves being vanishingly small in that area, resulting in a Fisher Information error that tends to extremely large values where the Bayesian decoder error is bounded. This regime disrespects the Kramer-Rao bound. **B:** Same as in A, but for Fig.1B. **C:** and **D:** Some quadruplets display a Fisher information that, due to their particular configuration of tuning curve slopes, tend to zero in certain points. Consequently the associated decoding error estimates grow to values which egregiously exceed the Bayesian decoding error. **E:** Spearman Rank correlation coefficients between the Bayesian decoding error and either $e^{-SSI(\theta)}$ (magenta) or $1/\sqrt{FI(\theta)}$ (orange). We can see that for $N = 648$ quadruplets the Sensitivity error estimate correlates with the Bayesian error more than the Fisher information error does. **F:** Difference between Sensitivity and Fisher performance in terms of Spearman Rank coefficients of correlation to the Bayesian decoding error. The differences are done quadruplet-by-quadruplet, showing that for all of them the Sensitivity performance correlates with the Bayesian error more than the Fisher performance does.

A system which is highly sensitive to a particular stimulus should be expected to yield a small decoding error when probed for that stimulus. The Stimulus specific information ($SSI$) measures the average information carried by responses conditioned on stimuli that evoked them, so we should expect it to bear some relation to the decoding error. More precisely, upon stimulation a system generates a response sampled from a likelihood distribution conditional on the specific stimulus presented ($p(\mathbf{r}|\theta)$); this response should "point" to the stimulus which causes it, and the degree to which a one-to-one mapping can be established corresponds to the sensitivity of the system to the stimulus in question.

In order to verify that the SSI has those desired properties, and is therefore a good measure of sensitivity, we need to link it with the error of a Bayesian decoder. Up to higher order corrections, and provided that the posterior $p(\theta|\mathbf{r})$ has a small enough variance, we can ignore the fact that the stimulus ensemble is bounded, and use the fact that the Gaussian distribution maximizes the entropy of a random variable to obtain an approximate lower bound of the Bayesian decoding error as a function of the SSI:

$$\sigma^2_{dec} \gtrapprox \sigma^2_{SSI}(\theta) \equiv \frac{|\Theta|^2}{2\pi e} e^{-2 \cdot SSI(\theta)} \ , \tag{6}$$

where $\sigma^2_{dec}$ is the decoding error. Eq. (6) proves that the SSI can provide an estimate of the decoding error, and so can be used to quantify the stimulus sensitivity.

The Fisher information is another quantity from information theory that can be linked with decoding error via the Kramer-Rao bound [43]. For our system, neglecting noise correlations and therefore supposing conditionally independent cell responses, the Fisher information takes on a simple form dependent only on the properties of the tuning curves constituting the quadruplet:

$$FI(\theta) = \sum_{k=1}^{4} \frac{\left(f'_{TC_k}(\theta)\right)^2}{f_{TC_k}(\theta)} = \sum_{k=1}^{4} FI_k(\theta) \tag{7}$$

Simplicity and relatively direct interpretability make the Fisher information an often-used measure of performance in neuroscience [42, 44, 45]. That utility notwithstanding, the Fisher information presents a striking vulnerability in the specific case of small populations with tuning curves that can flatten in extended regions of the stimulus ensemble, such as the Flat-Topped Von Mises. In such a case, the probability that all the tuning curve slopes (almost) equal zero is non-negligible (FigS1, A-D, top panels). Consequently, the Fisher information will take on small values in those regions (FigS1, A-D, middle panels), which will in turn lead to large values of its inverse - the Fisher information decoding error estimate from the Cramer-Rao bound (FigS1, A-D, bottom panels). This would lead to the implausible conclusion that the region is uninformative, even if there a cell is spiking maximally.

Fig.s S1-A to E show that $\sigma^2_{SSI}(\theta)$, from Eq. (6), correlates with the Bayesian decoding error more strongly than does the Fisher error. The medians of the histograms are respectively 0.77 and 0.52, while the $mean \pm std$ are, again respectively, $0.71 \pm 0.24$ and $0.47 \pm 0.37$. Furthermore, there is a substantial number of quadruplets for which the Fisher error negatively correlates with the Bayesian error. On a quadruplet-by-quadruplet basis the sensitivity error more closely correlates with the Bayesian decoding error compared to its Fisher information counterpart (FigS1-F).

In summary, the Fisher information is a commonly used and often powerful tool for estimating a lower bound on the decoding error, and could in principle provide a robust measure of sensitivity. However in specific regions where the tuning curves flatten, the Fisher information fails to provide a plausible decoding error estimate. On the other hand, the sensitivity error based on SSI does not suffer from such a drawback. Instead it closely approximates the Bayesian decoding in the small noise limit. Finally, its high correlation with the Bayesian decoding error is a property which one should hope to find in an appropriately defined measure of sensitivity.
